# All-optical voltage imaging-guided postsynaptic single-cell transcriptome profiling with Voltage-Seq

**DOI:** 10.1101/2023.11.24.568588

**Authors:** Veronika Csillag, JC Noble, Daniela Calvigioni, Björn Reinius, János Fuzik

**Affiliations:** Department of Neuroscience, Karolinska Institutet, Stockholm, Sweden; Department of Medical Biochemistry and Biophysics, Karolinska Institutet, Stockholm, Sweden

## Abstract

Neuronal pathways recruit large postsynaptic populations and maintain connections via distinct postsynaptic response types (PRTs). Until recently, PRTs were only accessible as a selection criterion for single-cell RNA-sequencing (scRNA-seq) through probing by low-throughput whole-cell electrophysiology. To overcome these limitations and target neurons based on specific PRTs for soma collection and subsequent scRNA-seq, we developed Voltage-Seq. An on-site analysis tool, VoltView, was created to guide soma harvesting of specific PRTs using a classifier based on a previously acquired connectome database from multiple animals.

Here, we present a detailed step-by-step protocol, including setting up the optical path, the imaging setup, detailing the imaging procedure, and analysis, the reagents and protocols, complete procedure for sequencing library preparation, and provide other tips and pitfalls to assist researchers in conducting high-throughput all-optical voltage imaging, and to obtain the single-cell transcriptomic data from selected postsynaptic neurons.

Voltage-seq workflow can be completed in ∼6 weeks including 4-5 weeks of viral expression of the Voltron sensor. The approach allows researchers to resolve the connectivity ratio of a specific pathway and explore the diversity of PRTs within that connectome. Furthermore, the high throughput in conjunction with quick analysis gives unique access to find specific connections within a large postsynaptic connectome. Voltage-seq also allows the investigation of correlations between connectivity and gene expression changes in a postsynaptic cell-type-specific manner in both excitatory and inhibitory connections.

## Introduction

Neuroscientists aim to develop a more comprehensive understanding of the nervous system. This effort robustly benefits from complex and high-throughput investigation of neuronal circuits. Multimodal identity of neurons became accessible with Patch-Seq^1,2^, where intrinsic electrophysiology, morphology and gene expression profile could be resolved within the same investigated neuron. Neuronal connectivity testing requires the detection of subthreshold postsynaptic excitatory (EPSP) and inhibitory (IPSP) responses. EPSPs and IPSPs have so far only been detected by classical intracellular whole-cell recordings. Probing postsynaptic connectivity with the combination of whole-cell recordings and optogenetic stimulation of incoming axon terminals had been performed for almost a decade^3^. However, this approach came with the inherently low throughput of whole-cell recordings of 10-15 neurons from a recording day and a skilled person. Furthermore, to find specific PRTs without a preview of the postsynaptic responses and obtain the molecular profile of a certain postsynaptic neuron is excessively challenging.

To address these limitations, we developed a technique termed Voltage-Seq. First, we optimized *ex vivo* all-optical voltage imaging using the Voltron sensor^4^. Genetically encoded fluorescent voltage indicators (GEVIs) report subthreshold voltage changes of both polarities^5,6^. GEVIs can thus capture both optical EPSPs (o-EPSPs) and optical IPSPs (o-IPSPs) and can detect single action potentials (APs). Harnessing the potential of the GEVIs fine sensitivity at this high throughput, we could simultaneously probe dozens of connections of up to 1000-1500 postsynaptic neurons on a recording day. The high-throughput all-optical connectivity testing generates a large data stream. To identify PRTs exhibiting particular characteristics (such as bursting or o-IPSPs) during the experiment, it is essential to rapidly analyze the voltage imaging data on-site, ideally within minutes. For this purpose, we built VoltView 1.0, capable of analyzing a field of view (FOV) with 50-60 neurons imaged at 600 Hz with 7 consecutive sweeps in about 1 minute. This rapid analysis gave access to combine a high-throughput connectivity screening with the single-cell molecular profiling of selected neurons of the postsynaptic connectome. We created Voltage-Seq, that used on-site analysis with VoltView to locate postsynaptic neurons with specific PRTs and to navigate soma harvesting to obtain the RNA transcriptome of neurons with target PRTs. We used the neuronal pathway from the ventromedial hypothalamus (VMH) to the periaqueductal gray matter (PAG) as a model to optimize the Voltage-Seq methodology, to resolve PRTs in the entire VMH-PAG connectome with all-optical voltage imaging, and to select specific neurons for somatic harvesting and subsequent scRNA-seq. We challenged Voltage-Seq to locate sparse GABAergic neurons in the VMH-PAG connectome guided by the on-site analysis in VoltView.

This protocol provides comprehensive guidance for implementing Voltage-Seq from the establishment of all-optical voltage imaging with the Voltron sensor, detailing the setup parts list and blueprint, the optical path, software use and analysis output. We also describe the workflow of imaging, analyzing, and harvesting with successful and failed examples. Importantly, we explain and illustrate the possible light artefacts that could influence the quality of all-optical imaging. We detail the complete sequencing library preparation protocol and the routine to get single-cell gene expression data. Within the procedural section, specific attention is given to pitfalls and the details of troubleshooting to ensure a successful application of our Voltage-Seq method.

### Overview of the procedure

The Voltage-seq workflow takes approximately 5-6 weeks, with 4-5 weeks for the optimal viral expression of the Voltron sensor, 1 day of data and single-cell soma collection, 2-3 days of scRNA-seq of a batch (∼100 neurons) and a few days of post-hoc and optical physiology (o-phys) analysis, classification, and anatomical mapping.

The process starts with the viral expression of somatic targeting Voltron (Voltron-ST) in the chosen postsynaptic areas and the expression of Channelrhodopsin-2 (ChR2) in the presynaptic neuronal pathway. In our experience, 4-5 weeks of expression time for Voltron-ST resulted in a satisfactory signal-to-noise ratio, while at 2-3 weeks of expression we observed weak or no signal. ChR2 expression becomes sufficiently high after 2-3 weeks. All-optical data collection and neuronal harvesting for Voltage-Seq from an animal has a time window of 7-8 hours which is limited by the e*x vivo* survival of brain slices. The voltage imaging data acquisition takes 1 minute per FOV, followed by the on-site analysis with VoltView, requiring 1-1.5 minutes per FOV. The process of soma harvesting with first locating the neuron with the desired PRT, harvesting, and transferring to the tube with lysis buffer (LB) takes ∼15-20 minutes per neuron. Optionally, harvested soma samples can be accumulated at -80°C throughout months to proceed to sequencing library preparation with larger batches (100-200 neurons) of samples. The sequencing workflow includes quality control of cDNA of the single neuronal samples after reverse transcription (RT), also upon library preparation, and after scRNA-seq. In the post-hoc analysis phase, the recorded data can be analyzed in further detail, facilitating the classification with hierarchical clustering and 3D mapping of neuronal PRT types and clusters.

### Imaging setup and imaging protocol

One of the most critical aspects of our methodology is to minimize the cross activation of ChR2 by the yellow light used to excite the Voltron sensor. We used 473 nm opto-stimulation for ChR2 activation, while simultaneously imaged with 585 nm excitation of the Voltron-ST labeled with Janelia Fluor dye-585 (JF585) (Fig. 2a). Interestingly, we experienced ChR2 cross activation resulting in a ∼5-6 mV somatic response with an earlier tested double-band excitation filter (59022x, Chroma) that transmitted excitation light between 450-490 nm and 550-590 nm. To well-separate the 585 nm light and prevent cross activation of ChR2, we used a dual-band excitation filter (59010x, Chroma) that transmitted excitation light between 475-500 nm and 570-595 nm (Fig. 2b). This resulted in a negligible ∼0.5-0.7 mV response measured on ChR2-expressing somas, but we could not induce synaptic release with this optical setup. This is essential for the clearance of all-optical imaging experiments. The 585 nm light was delivered at an excitation intensity of ∼14 mW/mm2, while the 473 nm light had an intensity of ∼2.5 mW/mm2 at the plane of imaging. Efficient segregation of emitted light from the excitation light was achieved by a band-pass emission filter (ET645/75, Chroma) and a dichroic mirror (T612lprx, Chroma) (Fig. 2b).

**Fig 1.**
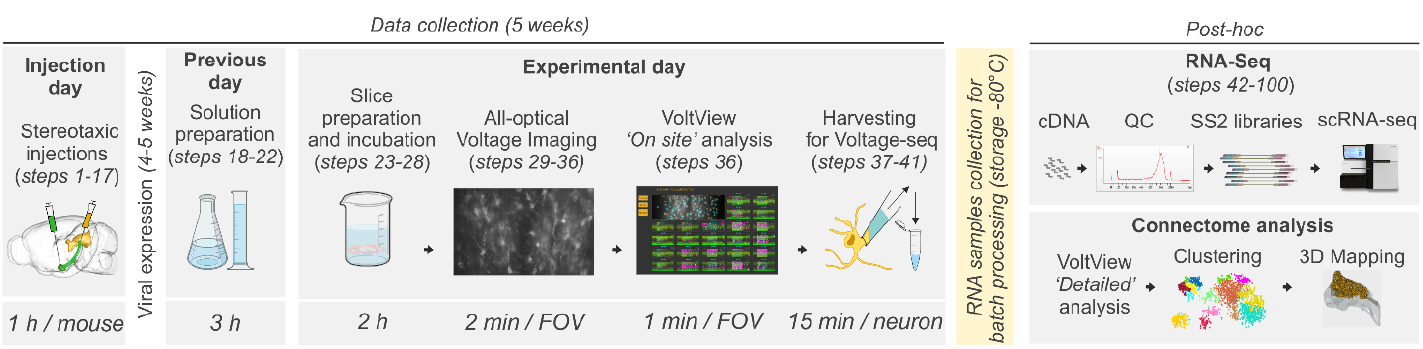
Experimental overview: Schematic summary outlining the sequence of experimental procedures. Each procedure is labeled with a step number and the name of the specific procedure, depicted above the respective illustrations. The length of each procedure is indicated at the bottom of the illustrations. The main experimental path contains steps from 1-100, alternatively followed by post-hoc connectome analysis.

**Fig 2.**
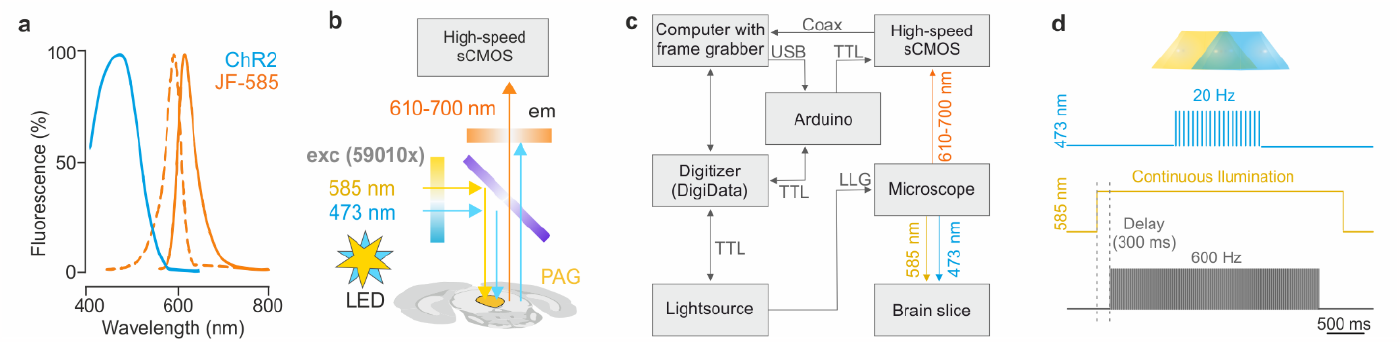
All-optical imaging setup and optics. **a**, Excitation, and emission profile of ChR2 (blue) and JF-585 (orange). **b**, Optical paths of all-optical voltage imaging *ex vivo* with double band excitation filter 59010x (blue to yellow), dichroic mirror T612lprx (purple), pass emission filter ET645/75 (orange) **c**, Block diagram of the connected instruments of the all-optical imaging setup. d, All-optical voltage imaging protocol of one typical sweep. (Top) TTL signal from DigiData activating the 473 nm channel of the light source to deliver 20 Hz opto-stimulation, (Middle) TTL signal from DigiData activating the 585 nm channel of the light source to continuously excite Voltron-JF585. (Bottom) TTL signal from Arduino at 600 Hz to trigger the acquisition of each frame.

During the all-optical voltage imaging experiments the computer coordinated all the instruments by sending commands via the analogue-digital converter (DigiData 1550B) (Fig. 2c). For high-speed voltage imaging we use a scientific Complementary Metal-Oxide-Semiconductor (sCMOS) camera (ORCA-Fusion BT, Hamamatsu). This camera ran the acquisition and saved the video files under the control of the HCImageLive software and received the frame triggers from an Arduino (UNO). The Axon patch-clamp amplifier (Multiclamp 700B) is reliable at patch-clamp experiments but cannot deliver high frequency TTL commands with accurate timing. The use of Arduinos is a viable alternative to create any frequency with high temporal accuracy. The Arduino was triggered by the DigiData to deliver 600 Hz frame triggers to the sCMOS following a protocol scripted and stored in it. Importantly, the computer was equipped with a frame grabber card (FireBird Dual CXP-6 PE8 G2 board v3AB) for high-speed data acquisition through coax cables (CoaXPress), only slower data acquisition is possible via an USB connection. The LED light source (Spectra-X, Lumencore) delivered light via a liquid light guide (LLG) and the color channels for voltage imaging and optogenetic stimulation were temporally coordinated by the DigiData, to have them in synchrony with the camera frame triggers. We created the protocol files for imaging in pClamp v10.4. that controlled the DigiData outputs, that also allowed to save the used protocols to track back the adjustments of settings during experiments (Fig. 2c). In our experimental setup, the optical stimulation and voltage imaging light are delivered through the objective lenses with the aperture of the fluorescent light path minimized to the area of the FOV. This adjustment serves the dual purpose of minimizing potential bleaching in the currently not imaged areas caused by the 585 nm light and restricts the opto-stimulation to the area of the currently imaged FOV.

The Voltron sensor is a Förster resonance energy transfer (FRET) protein which during imaging is continuously charged by a flow of power delivered by the imaging 585 nm light. Activation of the 585 nm light results in a sudden positive increase in the fluorescence of the Voltron in the illuminated area. To avoid recording this huge exponential light artefact, the protocol file commanding the light source and the camera initiates with the activation of the 585 nm imaging light and with a delay of 300 ms, the Arduino starts to send the 600 Hz frame triggers to the sCMOS camera (Fig. 2d). The protocol file consists of sweeps with 0.5 sec baseline, 1 sec opto-stimulation at 20 Hz with 473 nm blue light, and 1 sec post-stimulus period. Sweeps were separated by 7 sec inter-sweep intervals to give time for the neurotransmitter reuptake in the synaptic terminals and avoid synaptic depletion. We recorded at least 4 sweeps to average the extracted parameters for better classification. We chose 20 Hz frequency as opto-stimulation and only delivered the stimulus for 1 sec to identify connections while avoiding long-term synaptic potentiation.

This accurate orchestration of instruments, the appropriate separation of excitation wavelength of optogenetic actuator and the excitation wavelength of the Voltage sensor are essential steps in establishing controlled all-optical experiments. Bypassing the possible artefacts of the voltage sensor and calibrating the protocol to avoid excess bleaching is important to capture high-quality all-optical voltage imaging data.

### VoltView

We created VoltView to get a quick analysis of neuronal activity following all-optical voltage imaging. VoltView allows to choose neurons based on their PRTs, locate them in the FOV and approach them for soma harvesting. The On-site analysis was written in MATLAB (MathWorks) using custom-written scripts. It is specifically designed for the file structure generated by HCImageLive that creates the videos by concatenating the sweeps in one imaging session. Similarly generated file structures with another format such as “.TIF” would potentially be analyzed. During all-optical Voltron-imaging, Blue-spike artefacts are created by the 473 nm light pulses of the opto-stimulation, as they serve as additional energy injections into the continuously illuminated and excited FRET sensor (Fig. 3). In other words, during Voltron-imaging, the blue light pulses overcharge the Voltron and induce large spikes with opposing polarity to APs. The 1.0 version of VoltView was specifically designed for all-optical voltage imaging analysis with the Voltron sensor, hence it uses the Blue-spike artefacts of the Voltron-ST to detect the time points of optogenetic stimulations. This way the analysis adapts to on-site changes made in the command protocol, such as starting time, length, or frequency of optogenetic stimulation. dF/F of blue spikes are more than a magnitude larger in amplitude than that of neuronal signals. The extremely low expression level of Voltron-ST results in undetectable Blue-spikes (Fig. 3) and the On-site analysis will not run. Besides the analysis of single videos, multiple video comparison mode has interactive settings on-site to switch between parameters to define the color coding of increase or decrease of the chosen parameter in each imaged neuron. VoltView has Cellpose^7^ built-in, which is a highly trained and well-optimized generalist segmentation package that recognizes neuronal somas with high precision. The precise segmentation on the locally equalized and contrast enhanced FOV images enabled us to use Concentric analysis, defining “Inner” and “Outer” areas to clean the data from single photon background by discarding neurons that have smaller signal on their somas compared to surrounding “Inner” and “Outer” concentric areas.

**Fig 3.**
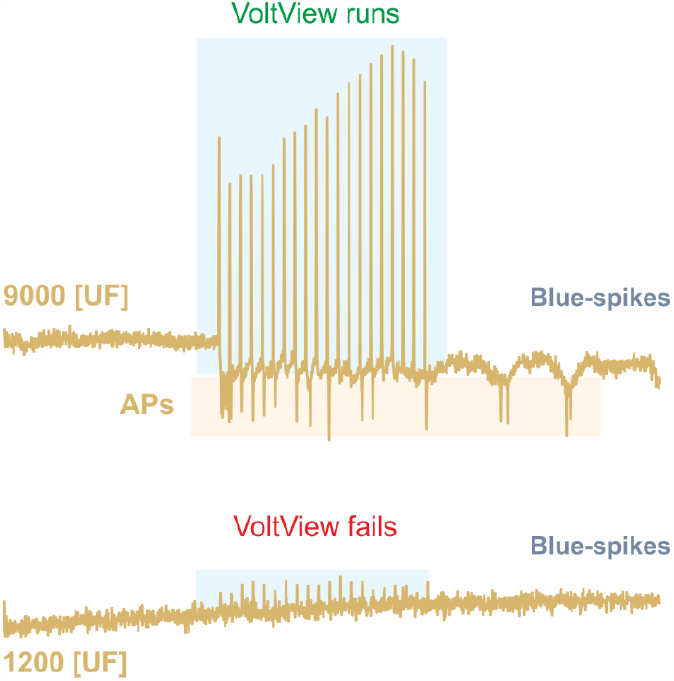
Blue-spike artefacts. Blue-spikes evoked by 20 Hz opto-stim of an all-optical imaged neuron (blue rectangles) that are detected by VoltView; Blue-spikes have reversed polarity to APs (top). Blue-spikes are small to be reliably detected in a neuron with low Voltron-ST expression (bottom).

Importantly, VoltView not only filters out traces that can be visually explored but also detects and extracts various o-phys parameters such as APs, EPSPs, IPSPs, and Bursts. It provides 29 different o-phys parameters for the classification of PRTs (Table 3.). VoltView features a built-in classifier that can be updated after each experiment, continuously improving the accuracy of PRT classification by incorporating more o-phys data into the clustering analysis. These updates generate cluster centroids used by VoltView for on-site classification, such data has to be copied into the folder of VoltView and named “VMH_PAG_Cluster_centroids” or a user can search for this variable in the script accessible online (https://zenodo.org/record/8030176) and rename it to the desired variable name.

**Table 1:**
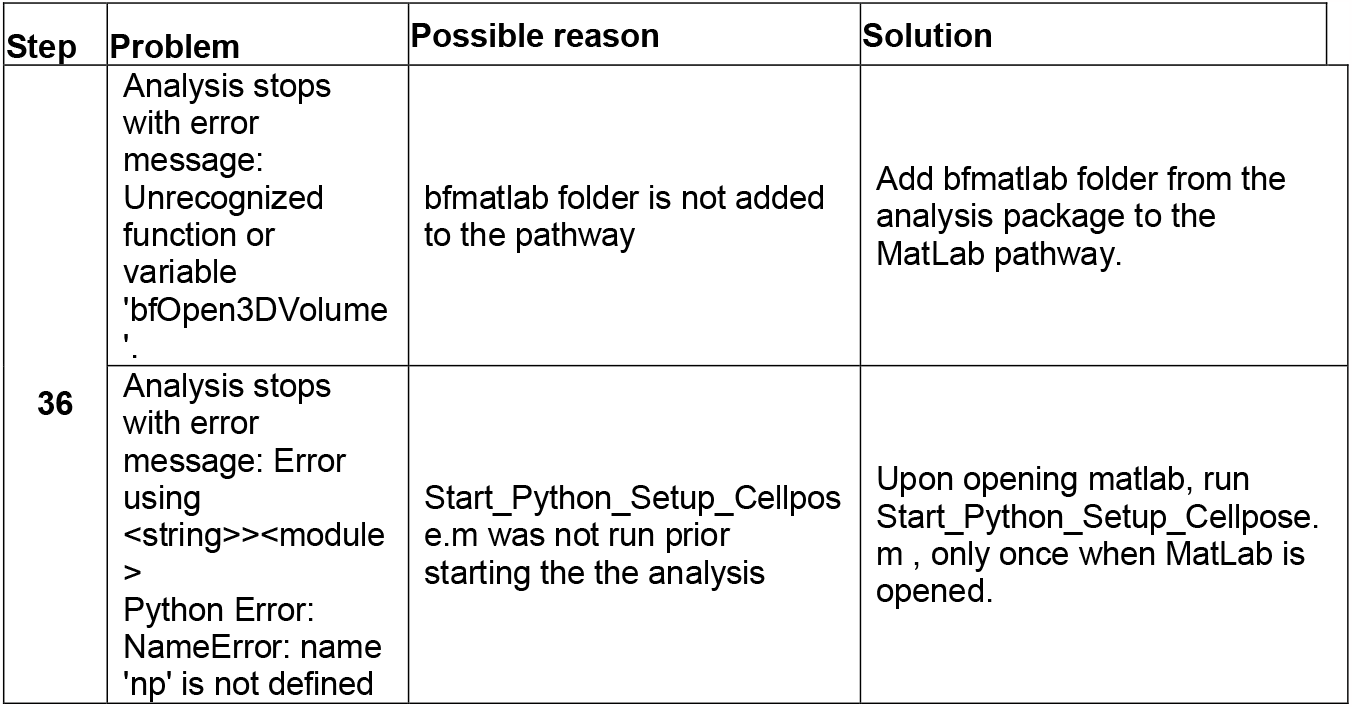
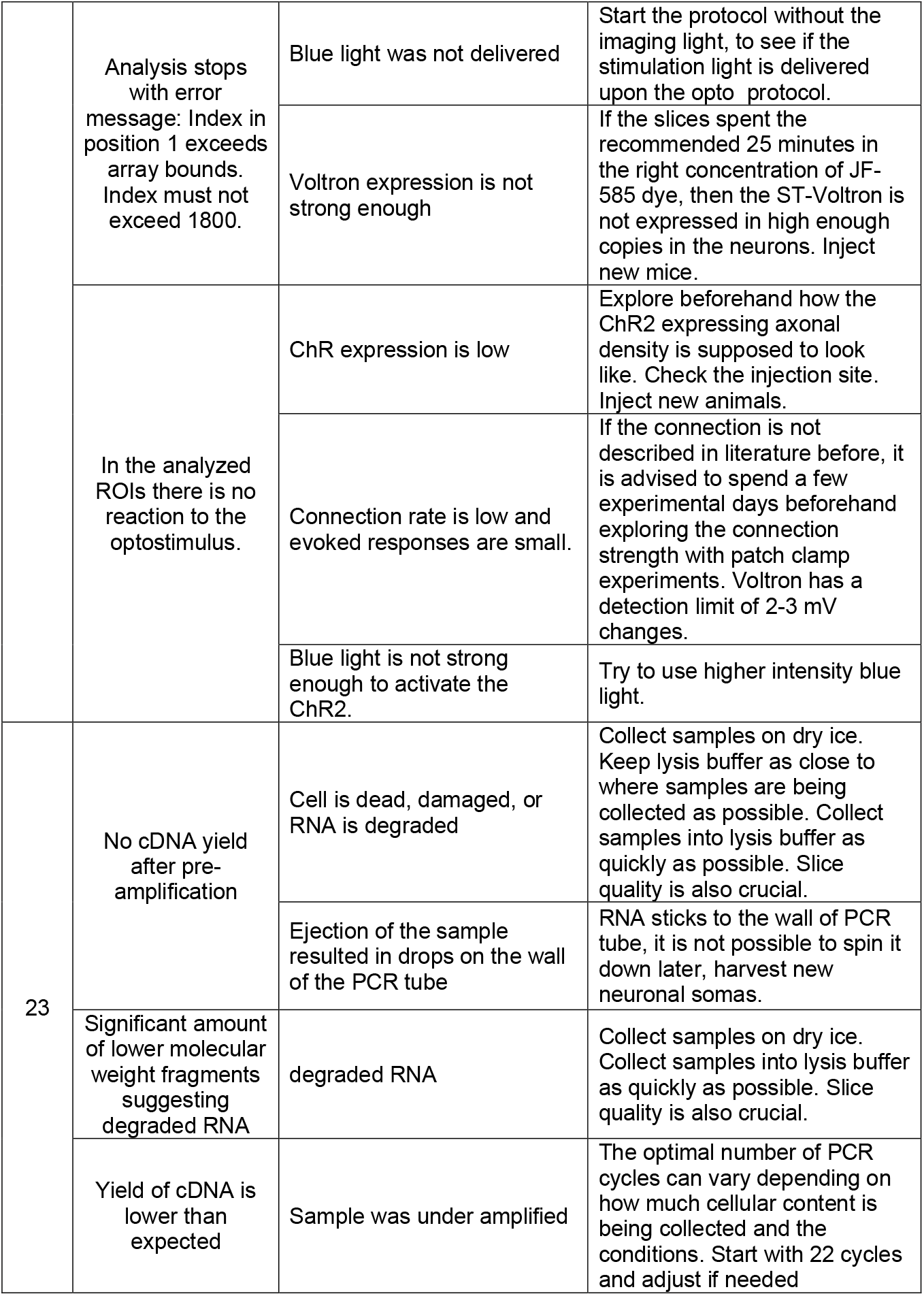
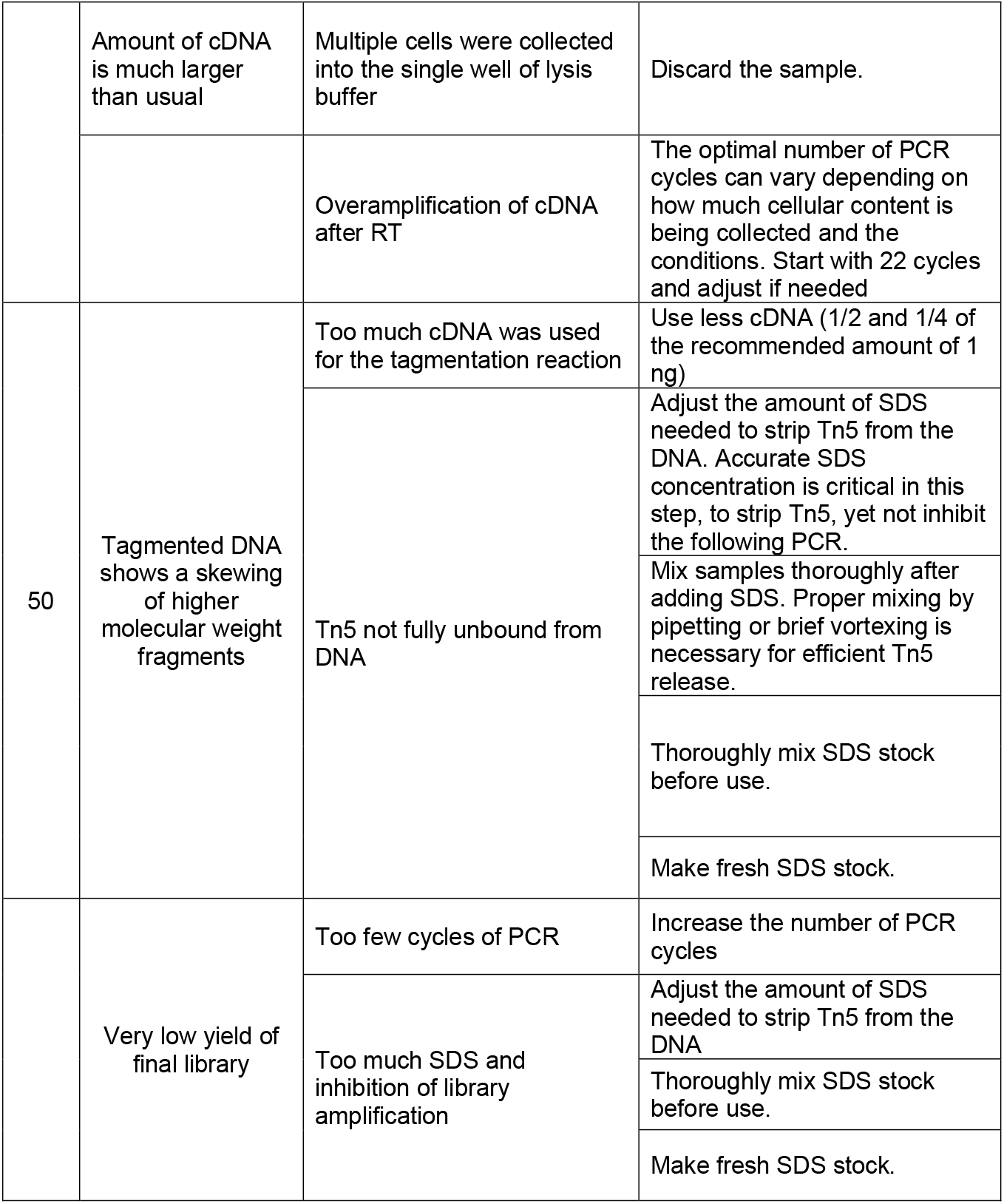
Troubleshooting.

**Table 2:**
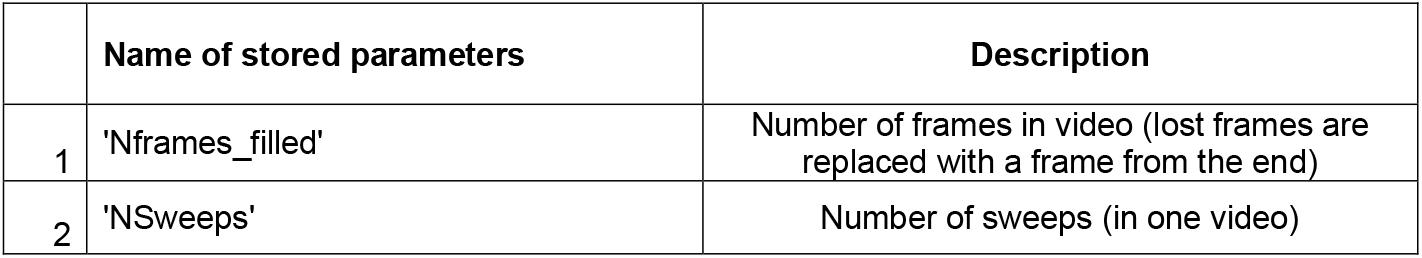
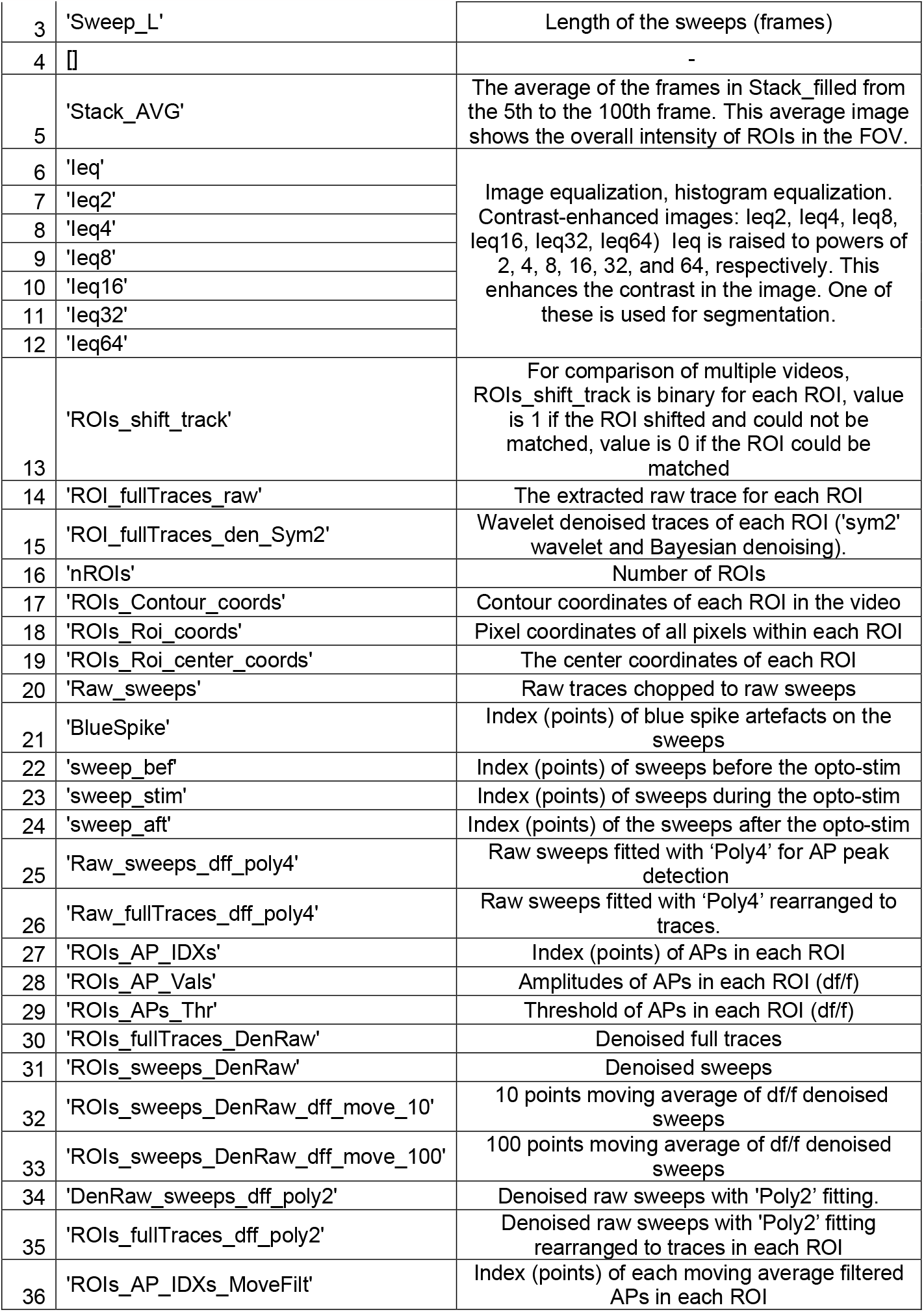
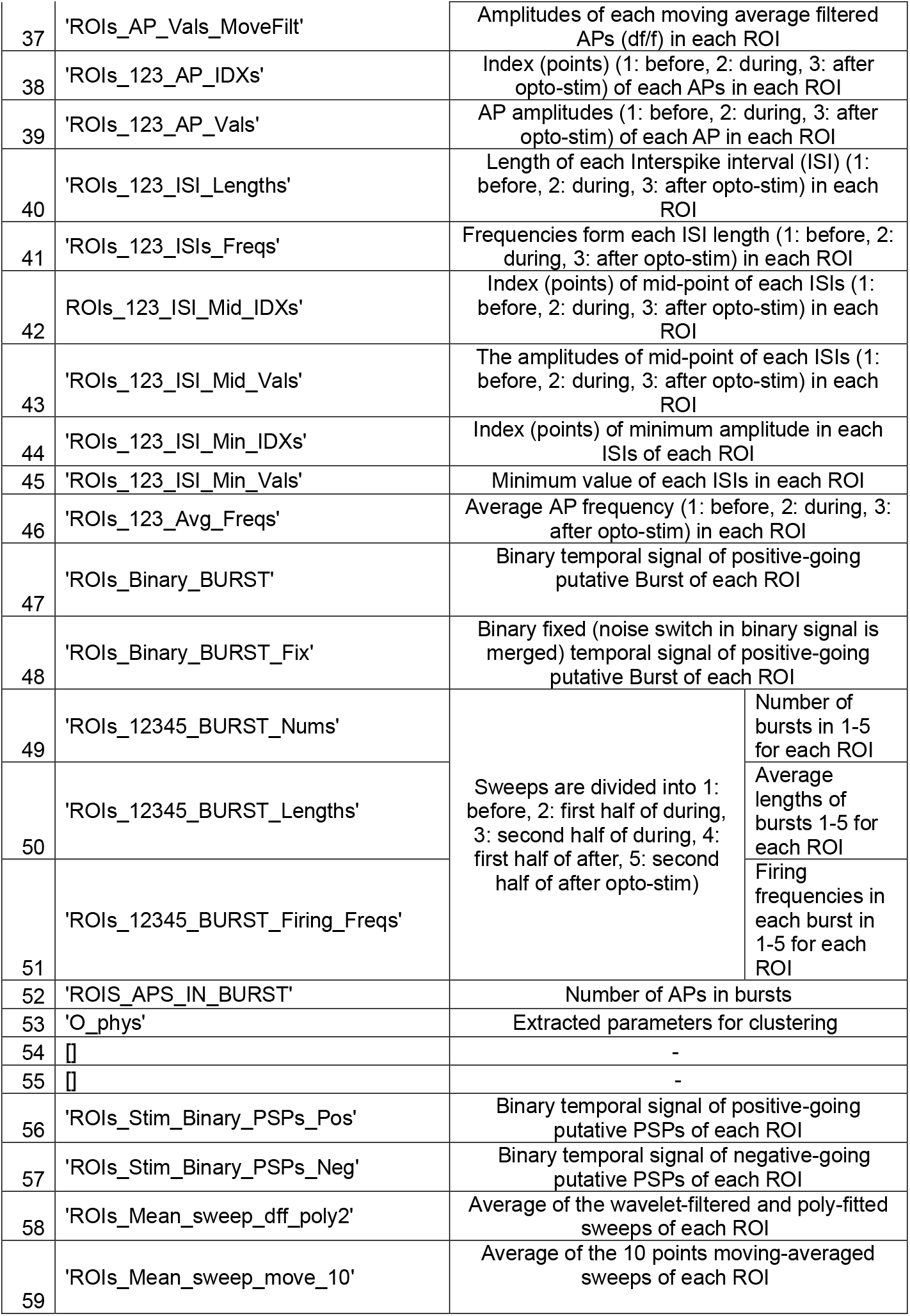
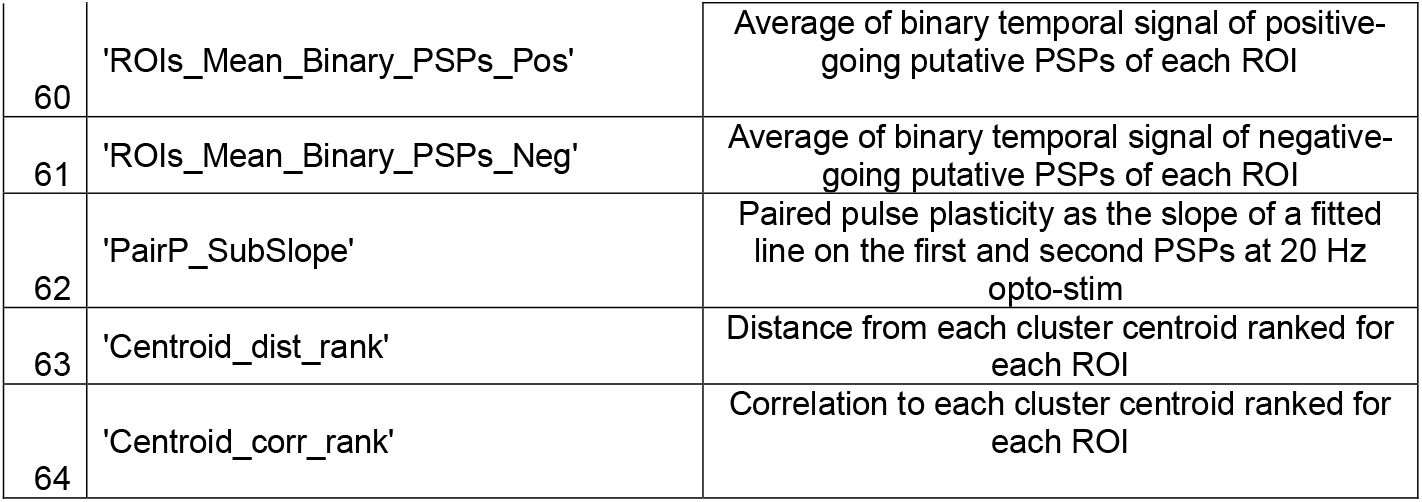
Extracted raw and processed o-phys signals.

**Table 3.** Extracted o-phys parameters.

|  |  |  |
| --- | --- | --- |
| O_phys (1) | Avg_Freq_bef | Average frequency of o-APs detected before 20 Hz opto-stim in all 6-7 sweeps |
| O_phys (2) | AP_num_bef | total number of o-APs detected before 20 Hz opto-stim in all 6-7 sweeps |
| O_phys (3) | PSP_pos | Number of o-EPSPs during the 20 Hz opto-stim. |
| O_phys (4) | PSP_neg | Number of o-IPSPs during the 20 Hz opto-stim. |
| O_phys (5) | PP1 | Paired pulse ratio of the second o-PSP amplitude compared to the first o-PSP amplitude. |
| O_phys (6) | PP2 | Paired pulse ratio of the third o-PSP amplitude compared to the second o-PSP amplitude. |
| O_phys (7) | SSlope_21 | 10 point moving average of opto-stim is divided into 4 x 250 ms quarters (q1-q4), SSlope_21 = q2/q1 |
| O_phys (8) | SSlope_all | SSlope_all = q4/q1 |
| O_phys (9) | Burst_Freqs_stim | Firing rate inside detected bursts during 20 Hz opto-stim. |
| O_phys (10) | Burst_Freqs_aft | Firing rate inside detected bursts after 20 Hz opto-stim. |
| O_phys (11) | Burst_Aps_perc | "AP % in burst" quantified the number of o-APs inside burst periods. |
| O_phys (12) | perc_out | Percentage of APs outside detected bursts. |
| O_phys (13) | AP_Onset_stim | Delay of the first o-AP during 20 Hz opto-stim. |
| O_phys (14) | AP_Freq_stim_H1 | AP freq average firing rate on the first half (H1) of during the 20 Hz opto-stim. |
| O_phys (15) | AP_Freq_stim_H2 | AP freq average firing rate on the second half (H2) of during the 20 Hz opto-stim. |
| O_phys (16) | AP_Freq_stim_H2_H1 | AP freq change of firing rate during opto-stim comparing the second half (H2) with the first half (H1) |
| O_phys (17) | AP_Freq_aft_H1 | AP freq average firing rate on the first half (H1) of after the 20 Hz opto-stim part. |
| O_phys (18) | AP_Freq_aft_H2 | AP freq average firing rate on the second half (H2) of after the 20 Hz opto-stim part. |
| O_phys (19) | AP_Freq_aft_H2_H1 | AP freq 2nd/1st (Post)" quantified the change of firing rate after the 20 Hz optostimulus comparing the second half (Post-2) with the first half (Post-1). |
| O_phys (20) | BIMOD_Binary | 1 is bimodal (when coefficient was above 5/9) and 0 is non-bimodal o-AP peak distribution |
| O_phys (21) | BIMOD_Coeff | Calculated with Sarle's bimodality coefficient as the square of skewness divided by the kurtosis, value for uniform distribution is 5/9, values greater than that indicate bimodal (or multimodal) distribution. |
| O_phys (22) | Burst_length_stim_h1 | Average length of burst periods during the first half(H1) of the 20 Hz opto-stim period. |
| O_phys (23) | Burst_length_stim_h2 | Average length of burst periods during the second half(H2) of the 20 Hz opto-stim period. |
| O_phys (24) | Burst_length_aft_h1 | Average length of burst periods during the first half(H1) of after the 20 Hz opto-stim period. |
| O_phys (25) | Burst_length_aft_h2 | Average length of burst periods during the second half(H2) of after the 20 Hz optostimulus period. |
| O_phys (26) | Burst_number_stim_h1 | Number of detected bursts in the first half(H1) of the 20 Hz opto-stim period. |
| O_phys (27) | Burst_number_stim_h2 | Number of detected bursts in the second half(H2) of the 20 Hz opto-stim period. |
| O_phys (28) | Burst_number_aft_h1 | Number of detected bursts in the first half(H1) of after the 20 Hz opto-stim period. |
| O_phys (29) | Burst_number_aft_h2 | Number of detected bursts in the second half(H2) of after the 20 Hz opto-stim period. |

### Limitations

We provide a detailed description of how to apply the Voltage-seq method in mice. Viral incubation times in other rodents might differ. We specifically probed synaptic connectivity in the VMH-PAG connectome, investigating other brain areas might require additional calibration for example: viral expression time, optical stimulation intensity and frequency. It may be necessary to assess the synaptic strength in the brain region of interest with preceding whole-cell patch-clamp recordings before implementing Voltron-imaging, as Voltron had ∼2-3mV detection limit in our hands.

While our original paper demonstrated negligible crosstalk with exciting the JF-585 ligand^8^, using JF-646 could further decrease crosstalk, although the reported dF/F values in response to comparable membrane voltage changes are substantially smaller. We think that the validation of the lack of crosstalk is always essential when implementing different combinations of GEVIs and optogenetic actuators. All-optical voltage imaging technique is currently suitable for investigating only one projection within one postsynaptic area, because of the overlapping excitation wavelength properties of GEVIs and opsins.

The imaging frequency is inversely proportional to the size of the FOV. With a 600 Hz frame rate, we were able to image a 200 um x 600 um area (that means 60-80 neurons on average). We chose this frequency to capture all action potentials and to observe disynaptic inhibition.

Our recorded data is saved as “.cxd” files, different files may need modification of the data loading step in VoltView. We also created a built-in classifier, which needs to be adjusted/rebuilt by the experimenters as it is built for VMH-PAG connectome.

While Voltage-seq offers approximately a two-times-higher throughput compared to classical Patch-seq experiments, it is still limited to approximately 20-25 neurons per day, of course these samples are already preselected based on connectivity analysis and specific PRT in response to a pathway-specific all-optical measurement.

## Materials

### Stereotaxic injections

#### Equipment

- Stereotaxic frame (Model 900/902, Harvard Apparatus)
- Positive displacement microcapillary pipettes (Drummond Scientific)
- Quintessential Stereotaxic Injector (Stoelting, Cat. No.: 53311)
- Puller for the microcapillary pipettes (any brand)
- Homotermic Blanket Control Unit (Harvard Instruments, Cat. No.: 50-7053)
- Drilling tip (Meisinger, HM71 Cat. No.: 2000071104004)
- Scissors (Dumont, Iris style)
- Forceps (Dumont, Dressing and Adson style)
- Hemostatic Forceps (any brand)
- Microdrill machine (Foredome, Cat. No.: K1070-21)

#### Consumables

- Mineral oil (Sigma Aldrich, Cat.No.:330779)
- Vaseline (any brand)
- Suturing material (Ethilon, Cat.No.:1696G)
- Qtips (any brand)
- Chlorhexidine (Fresenius Kabi, Cat.No.:537993)
- 70% Ethanol (any grade)
- 1 ml syringes and 31G needles (any brand)

#### Viral constructs

- pAAV-hsyn-flex-Voltron-ST (AAV1); Addgene, cat. no. 119036-AAV1; (at titer ≥2×10e12 viral genomes/ml)
- pAAV-hSyn-hChR2(H134R)-EYFP (AAV5); Addgene, cat. no. 26973-AAV5; (at titer ≥ 7×10e12 viral genomes/ml)
- pENN-AAV-hSyn-Cre-WPRE-hGH (AAV1); Addgene, cat. no. 05553-AAV1; (at titer ≥ 1×10e13 viral genomes/ml

## Brain slice preparation

### Equipment

- Transcardial perfusion tools, that is used only for electrophysiology or imaging experiments:
- scissors (Iris style), forceps
- 60ml syringe or perfusion pump
- Feeding needle (Agnthos, Cat.No.:7901)
- Vibratome (VT1200S, Leica, Germany)
- Thermo regulated water bath (VWR, Cat.No.:462-0555P)
- Superglue (Loctite Precision)
- Submerged incubation chamber for recovery and JF-dye incubation of brain slices (Custom built from 250 ml beaker and plastic mesh)

### Consumables

- 1 ml syringes and 31G needles (any brand)
- Blades for brain slicing (Gilette)

### Reagents

- KCl (Sigma Aldrich, Cat. No.: P9541)
- NaH2PO4 (AppliChem, Cat. No.: 131965)
- HEPES (Sigma Aldrich, Cat. No.: H4034)
- N-Methyl-D-Glucamin (NMDG) (Sigma Aldrich, Cat.No.:M2004)
- D-Glucose (Sigma Aldrich, Cat. No.: G7021)
- Hydrochloric Acid (Sigma Aldrich, Cat.No.:258148)
- NaHCO3 (Sigma Aldrich, Cat. No.: S6014)
- D-Sucrose (Sigma Aldrich, Cat. No.: S9378)
- L-ascorbic acid. (Sigma Aldrich, Cat. No.: A7631)
- CaCl2 (Cat. No.: C5080)
- MgSO4 (Cat. No.: M1880)
- Janelia Fluor 585 (Tocris, Cat. No.: 6418)

## Voltage-Seq – All-optical voltage imaging

### Equipment

- LED light source (Spectra X Light Engine, Lumencore, USA)
- Fast sCMOS camera (Orca Fusion-BT, Hamamatsu, Japan Cat.No.:C15440/20UP).
- PC with >32GB RAM of memory, with a Frame grabber card (Firebird CoaXPress Frame Grabber AS-FBD-2XCXP6-2PE8) to acquire the sCMOS data
- Axon DigiData 1550B (Molecular Devices, USA)
- Arduino Micro microcontroller (Arduino Uno)
- Filters and dichroic mirror for Voltage Imaging:
- dual-band excitation filter (Chroma, Cat.No.: 59010x) band-pass emission filter (Chroma, Cat.No.: ET645/75) dichroic mirror (Chroma, Cat.No.: T612lprx)
- Filter set for eYFP or GFP signal (Olympus, Cat. No.: N2713500)
- Objectives:
- Air objective, PLN 4X (Olympus, Cat. No.: 86-812)
- Water immersion objective, XLUMPLFLN20XW, (Olympus, Cat. No.: N2699600)
- Demagnifier 0.5x C-mount adapter (Olympus, U-TV0.5XCCN-2)
- Magnification changer to 2x (U-ECA, Olympus)
- DIC optics (Scientifica, condenser: OLY-38183; polarizer: OLY-38194; DIC prism: OLY-N2671800, analyzer: OLY-N2738400)
- SliceScope upright, fixed-stage microscope configured for epifluorescence illumination mounted on x-y stage (Scientifica, Cat. No.: S-SSMS-2000-00)
- Double PatchStar Micromanipulator (Scientifica, Cat. No.: S-PS-8300P)
- Holder for PCR tube for soma harvesting (custom made)
- Microforge to check harvesting capillary tip size (Narishige, MF-900)
- Pipette puller appropriate for fabrication of patch electrodes (e.g., Sutter Instruments, P series Flaming/Brown pipette puller; Narishige PC-100)

### Consumables

- Borosilicate glass capillaries, ID: 0.9 mm, OD 1.25 mm (Hilgenberg, Germany)
- Micro Loader tips (Eppendorf)

### Analysis with VoltView

Software dependencies:

- Camera software: HCImageLive software
- Code writing and testing was done in Windows 10:
- MATLAB 2021b
- Anaconda3 2021.11 (Python 3.9.7 64-bit)
- Cellpose 1.0

Hardware requirements:

- Minimum 8 GB memory but since files are large, 16 GB is advisory, 32GB is optimal.

### VoltView installation

1. Install Anaconda3 - installer version Anaconda3-2021.11-Windows-x86_64_3.9 (Typical install time on a “normal” desktop computer < 2 min)
2. Install Cellpose 1.0 (minimal version without GUI) from: (https://github.com/MouseLand/cellpose) Installation steps from the link: - Open an anaconda prompt / command prompt which has conda for python 3 in the path - Create a new environment with “conda create --name cellpose python=3.8” (We recommend python 3.8, but python 3.9 and 3.10 will likely work as well.) - To activate this new environment, run “conda activate cellpose” - To install the minimal version of Cellpose, run “python -m pip install cellpose” (Typical install time on a “normal” desktop computer < 2 min)
3. Install MATLAB 2021b (Typical install time on a “normal” desktop computer ∼30 min)
4. Download bioformats 6.11 package (OME - https://www.openmicroscopy.org/). and add “bfmatlab” folder to MATLAB path (in MATLAB: right click on folder/Add to path/Selected folders and subfolders)
5. Download “VoltView 1.0” (https://zenodo.org/record/8030176), copy to folder of choice

## Voltage-seq – SmartSeq2

### Equipment

- Centrifuge (Eppendorf, Centrifuge Cat. No.: 5810 R)
- Thermal cycler for PCR (Techtum Life Touch)
- DynaMag-PCR magnet for 0.2-ml PCR tubes (Thermo Scientific, Cat. No.: 492025)
- DynaMag-PCR magnet for 96-well PCR plates (Thermo Scientific, Cat.No.:12331D)
- DynaMag-2 magnet for microcentrifuge tubes (Thermo Scientific, Cat.No.:12321D)
- Qubit 2.0 fluorometer (Thermo Scientific, Cat. No.: Q32866)
- 2100 Bioanalyzer Instrument (Agilent, Cat. No.: G2939B)
- NextSeq 550 next-generation sequencing instrument (Illumina, Cat. No.: SY-415-1002)
- Microcentrifuge (VWR, Micro Star 17)
- Vortex (VWR, Analog Vortex Mixer)
- Plate Vortex Mixer (VWR, Advanced Plate Vortex Mixer)
- Pipettors (Gilson, Pipetman)
- Ice bucket

### Consumables

- Nonstick, RNase-free 1.5-ml microfuge tubes (Thermo Fisher Scientific, Cat.No.:AM12450)
- Sterile filter pipette tips 10, 30, 100, 200, and 1000 μl (Gilson, DIAMOND)
- Sterile serological pipettes 2, 5, 10, and 25 ml (Corning, Cat.No.:4486-4489)
- 8-strip, nuclease-free, 0.2-ml, thin-walled PCR tubes with caps (Sarstedt, Cat. No.: 72.991.002)
- 96-well PCR reaction plates (Thermo Scientific, Cat. No.: AB2596)
- 384-well PCR reaction plates (Thermo Scientific, Cat. No.: AB2596)
- Plastic film for PCR plates (Thermo Scientific, Cat. No.: AB0558)
- Conical tubes 15 ml (Sarstedt, Cat.No.:62.554.502)
- Conical tubes 50 ml (Sarstedt, Cat. No.: 62.559.001)
- 96-well black flat bottom plates (Greiner Bio-One, Cat.No.:655209)
- 384-well black flat bottom plates (Greiner Bio-One, Cat.No.:781209)
- Sterile disposable reagent reservoirs (Integra Biosciences, Cat. No.: 4331)
- Delicate task kim-wipes (KimTech Science by Kimberly Clark, Cat.No.:34133)
- NextSeq 500/550 High Output Kit v2.5 (150 Cycles) (Illumina, Cat.No.:20024907)

### Reagents

- Triton X-100 (100%) (Sigma, T8787)
- SEQURNA thermostable RNase inhibitor (50 U μl^−1^) (SEQURNA, SQ00201)
- dNTP Set (100 mM) (Invitrogen, 10297018)
- Ambion Nuclease Free Water (Ambion, AM9932)
- SuperScript™ II Reverse Transcriptase (Invitrogen,18064022). Comes with Superscript II first-strand buffer (5×) and DTT (100 mM)
- Betaine (5 M) (Sigma Aldrich, B0300)
- MgCl2 (1 M) (Invitrogen, AM9530G)
- KAPA HiFi HotStart ReadyMix, 2× (Roche, KK2601)
- 10x TAPS-Mg Buffer
- PEG 8000 (Sigma, P2139)
- KAPA HiFi HotStart plus dNTPs (Roche, KK2501)
- 20% SDS (Sigma 428018)
- Custom library index plates (custom Nextera index primers, IDT)
- Custom Tn5 (Picelli S. et al., 2014)
- 70% Ethanol (any grade)

### Oligonucleotide sequences

- Smart-seq2 Oligo dT - 5′-AAGCAGTGGTATCAACGCAGAGTACT30VN-3′
- Smart-seq2 TSO - 5′-AAGCAGTGGTATCAACGCAGAGTACATrGrG+G-3′
- ISPCR: 5′-AAGCAGTGGTATCAACGCAGAGT-3′

### Reagent Setup

- 10x TAPS-Mg Buffer – 100 mM TAPS, 50 mM MgCl2, pH 8.5 (adjusted with NaOH)
- 22% PEG Clean-up Beads – 1 mL of 1x TE-washed Sera-Mag Speed Beads washed and resuspended in 900 μl 1x TE, added to 49 mL of 22% PEG 8000, 10mM Tris pH 8.0, 1mM ETDA, 1M NaCl, 0.01% IGEPAL, 0.05% Sodium-Azide (equivalent to AMPure XP beads, Beckman Coulter).

### cDNA and Library Quality Control Reagents

- Qubit 1x dsDNA High-Sensitivity Assay Kit (Thermo Scientific, Q32854)
- Qubit Assay Tubes (Thermo Scientific, Q32856)
- High Sensitivity (HS) DNA Kit – Bioanalyzer Chips & Reagents (Agilent, 5067-4626)
- KAPA Library Quantification Kit for Illumina Platforms (KAPA Biosystems, KK4835)

### Timeline

Viral injection: 1 hour

Viral expression: 4-5 weeks

Solution preparation: 3-4 hours

Brain slice preparation and incubation: 1.5 hours

Incubation in JF dye: 25-30 minutes

All-optical voltage imaging and soma harvesting: 7-8 hours Library preparation: 1-2 days

ScRNA-Seq: 1 day

## Experimental steps

### Viral injections

1. Sterilize the working bench and tools with 70% ethanol.
2. Pull the glass injection pipette with the pipette puller to have the length to reach the targeted brain area and a sharp tip with a small diameter, to cause minimal damage to the brain.
3. To inject wild type animals, mix the pAAV-hsyn-flex-Voltron-ST (AAV1; Addgene) with pENN-AAV-hSyn-Cre-WPRE-hGH in 1:1 ratio. To inject Cre+ animals use the pAAV-hsyn-flex-Voltron-ST (AAV1; Addgene) alone.
4. Fill both capillaries with mineral oil and insert the microinjector. Push out a droplet of mineral oil at the end and fill the viral vectors from the front as follows:
5. Fill one injecting microcapillary with the mixture of the virus pAAV-hsyn-flex-Voltron-ST (AAV1; Addgene) with pENN-AAV-hSyn-Cre-WPRE-hGH in 1:1 ratio.
6. Prepare another capillary and fill it with pAAV-hSyn-hChR2(H134R)-EYFP (AAV5).

#### Suggestion

Inject the Voltron-expressing and Cre-expressing virus mixture and the hR2-expressing virus during the same surgery. The expression of ChR2 cannot be excessively high.

7. Anesthetize mice with isoflurane (2%) and place them into a stereotaxic frame (Harvard Apparatus, Holliston, MA). Before the first incision, administer analgesic Buprenorphine (0.1 mg/kg) and local analgesic Xylocaine/Lidocaine (4 mg/kg) subcutaneously. Body temperature of the mice must be maintained at 36°C with a feedback-controlled heating pad.

#### CRITICAL STEP

Test the animal”s reflexes to determine that a surgical level of anesthesia has been reached before proceeding with surgery.

8. Using hair trimmers, shave the scalp. Disinfect the skin with 70% ethanol using Qtips.

#### CRITICAL STEP

Only apply ethanol to the area where the incision is going to happen. Ethanol used on large areas of the skin on animals can contribute to body temperature loss, influencing the anesthesia.

9. Apply local anesthetics under the skin of the head, wait until the injected amount is absorbed.
10. Cut the skin with a single incision either with fresh scalpel blade or disinfected scissors.
11. Gently push aside connective tissue on top of the skull as needed for clear viewing. do the craniotomy at the targeted stereotaxic coordinates.

#### 12. CRITICAL STEP

The craniotomy procedure should proceed until the dura mater is still intact. The dura can be gently removed using sharp Dumont forceps or a needle. The dorso-ventral stereotaxic coordinates are relative to the brain surface. Any damage to the brain surface or excessive bleeding caused by the drill can alter the injection success rate.

13. For viral injections use micropipette attached on a Quintessential Stereotaxic Injector (Stoelting, Wood Dale, IL). Inject with the speed of 50-100 nl/minute.
14. Keep the injection pipette in place for 5 minutes after the injection. Slowly (100 μm/s) retract the pipette from the brain.
15. Clean the edges of the skin with Chlorhexidine and do the suturing to close the skin.
16. Give analgesics Carprofen (5 mg/kg) at the end of the surgery, followed by a second dose 18-24h after the surgery or as it is requested in the ethical permit.
17. Wait 4-5 weeks for viral expression.

## Solution preparation for brain slice *ex vivo* imaging

18. Make the following stock solutions, it takes 3-4 h the day before the all-optical voltage imaging experiment.
  - **NaHCO**_**3**_ **10x stock solution**

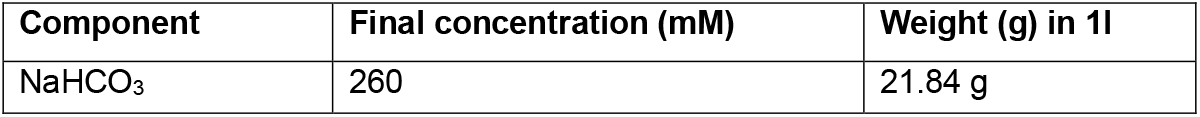
  - **D-Glucose 10x stock solution**

| Component | Final concentration (mM) | Weight (g) in 1l |
| --- | --- | --- |
| NaHCO<sub>3</sub> | 260 | 21.84 g |

  - **MgSO**_**4**_ **stock solution**

- **D-Glucose 10x stock solution**
| Component | Final concentration (mM) | Weight (g) in 1l |
| --- | --- | --- |
| D-Glucose | 200 | 36.02 |

  - **CaCl**_**2**_ **stock solution**

- **MgSO<sub>4</sub> stock solution**
| Component | Final concentration (M) | Weight (g) in 200 ml |
| --- | --- | --- |
| MgSO<sub>4</sub> | 1 | 49.3 |

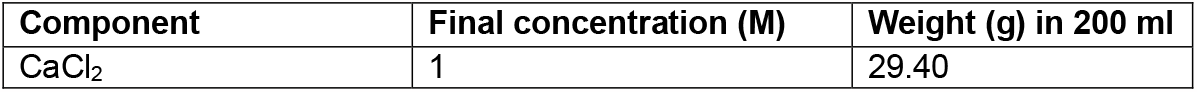
  - **Cutting/Recovery 10x stock**

- **CaCl<sub>2</sub> stock solution**
| Component | Final concentration (M) | Weight (g) in 200 ml |
| --- | --- | --- |
| CaCl<sub>2</sub> | 1 | 29.40 |

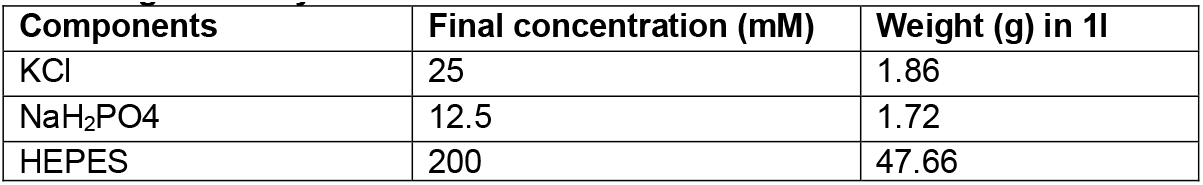
  - **aCSF 10x stock**

- **aCSF 10x stock**
| Component | Final concentration (mM) | Weight (g) in 1l |
| --- | --- | --- |
| NaCl | 124 | 72.46 |
| KCl | 25 | 1.86 |
| NaH<sub>2</sub>PO<sub>4</sub> | 1.25 | 1.72 |

19. **Using the stock solutions above, make the 1x solutions**.
  - **Sucrose containing cutting solution 1x (used in 2 days)**

  - **NMDG containing recovery solution 1x (used in 4 days)**

19. Using the stock solutions above, make the 1x solutions. - **Sucrose containing cutting solution 1x (used in 2 days)**
| Component | Final concentration (mM) | Amount in 500 ml |
| --- | --- | --- |
| NaCl | 80 | 2.33 g |
| Sucrose | 75 | 12.83 g |
| Cutting/Recovery 10x stock |  | 50 ml |
| D-Glucose 10x stock | 20 | 50 ml |
| NaHCO<sub>3</sub> 10x stock | 26 | 50 ml |

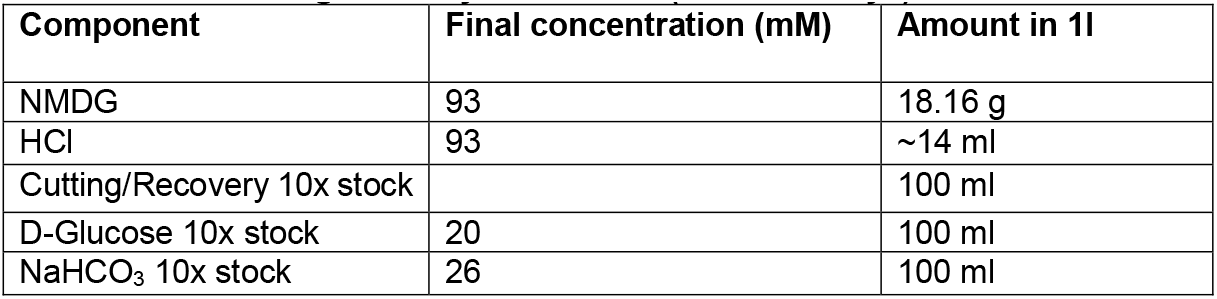 First take 500 ml DW and dissolve the NMDG, add utting/Recovery 10x stock k, D-Glucose 10x stock, NaHCO_3_ 10x stock and use 5M HCl to set the pH = 7.5.
  - **Sucrose-NMDG mixed solution 1:1 ratio (s-NMDG solution) – 1x**

  - **Recording aCSF (R-aCSF) - 1x**

| Component | Final concentration (mM) | Amount in 500 ml |
| --- | --- | --- |
| Sucrose containing cutting solution 1x |  | 250 ml |
| NMDG containing recovery solution 1x |  | 250 ml |
| CaCl<sub>2</sub> 1M stock | 0.5 | 250 µl |
| MgCl<sub>2</sub> 1M stock | 6 | 3 ml |
| Sodium-L-ascorbate | 3 | 1g |

  - **JF-dye incubating solution** Janelia Fluor 585 (Tocris, Cat. No.: 6418) JF-dye is dissolved in DMSO to a stock of 1 μM. Dilute 100 nmol JF-585 powder in 0.5 ml DMSO. Dilute it in 100 mL freshly made aCSF, aliquot it into 10 ml stocks. Freeze the 10ml aliquots at -20°C. Before the experiments, thaw and dilute aliquot to 15x with aCSF to a final volume of 150 ml to get 50 mM concentration.

- **Recording aCSF (R-aCSF) - 1x**
| Component | Final concentration (mM) | Amount in 2l |
| --- | --- | --- |
| NaHCO<sub>3</sub> 10x stock | 26 | 200 ml |
| D-Glucose 10x stock | 20 | 200 ml |
| R-aCSF 10x stock |  | 200 ml |
| CaCl<sub>2</sub> 1M stock | 2 | 4 ml |
| MgCl<sub>2</sub> 1M stock | 1 | 2 ml |

20. **Harvesting buffer in the harvesting capillary (HB):**
  - 90 mM KCl
  - 20 mM MgCl_2_ Pipette 4.5 ml 1M KCl stock and 1 ml 1M MgCl_2_ stock into 44.5 ml ultrapure distilled water.
21. **Prepare cell lysis buffer (LB):** One PCR tube contains 4 μl of LB

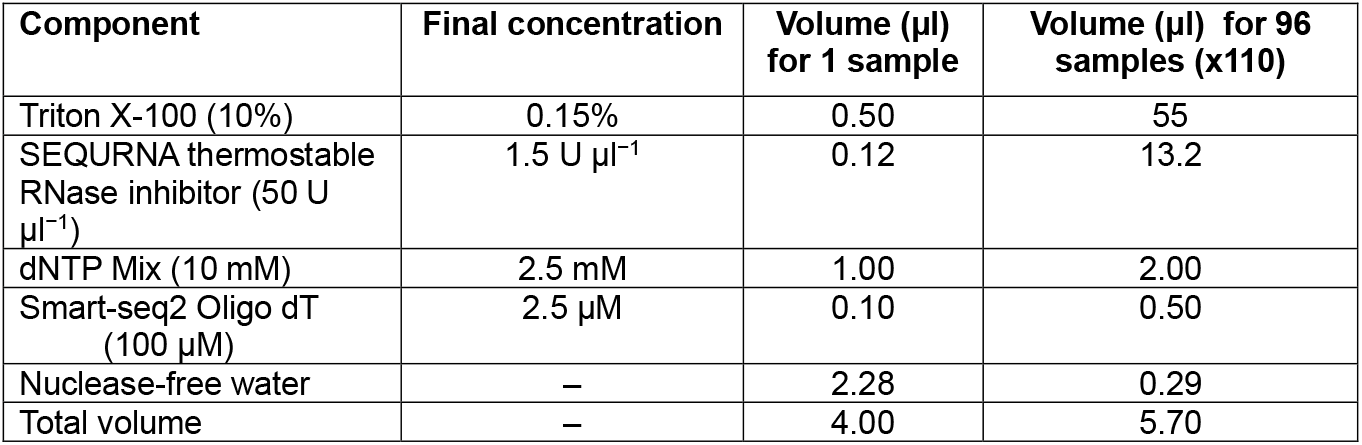

Premade PCR tubes with LB can be stored at −20°C or −80°C until use.

### Brain slice preparation *ex vivo*

Slice quality is crucial for voltage-imaging similarly to what is required for patch-clamp recordings. After 4-5 weeks of Voltron-ST expression, prepare brain slices following these steps:

22. Anesthetize mice with i.p. injection of 50 μl Na-pentobarbital (60 mg/kg) and transcardially perfuse with 4-8 °C S-NMDG solution^9^.

#### Suggestion

This step is optional, but beneficial as it improves slice quality and longevity by reducing excitotoxicity due to the removal of blood.

23. Carefully remove the brain after the transcardial perfusion from the skull within 1 minute (it is not necessary to do this in seconds or in a rush) and place it into 4-8 °C S-NMDG solution. Block the brain for the desired brain region and slicing angle and fill the buffer tray of the vibratome with the 4-8 °C S-NMDG solution and ensure continued carbogenation (95% O_2_ and 5% CO_2_) throughout the procedure.
24. Cut 250 μm thick coronal slices with the speed of 0.08 mm/sec in the 4-8 °C S-NMDG solution.

#### Suggestion

Slow slicing (0.03-0.05 mm/s) can improve slice surface quality.

25. Slice labeling: After finishing the cutting of a single brain slice, mark its orientation by cutting one corner with a surgical blade. Throughout the entire process, maintain this orientation for each slice. This practice ensures accurate mapping of the imaged neurons, since the distance of the imaged planes is going to be 250 μm.
26. After completion of the sectioning of each slice, collect them using a glass Pasteur pipette and transfer them into a pre-warmed (34 °C) recovery chamber filled with 150 ml of carbogenated S-NMDG solution. Every slice spends 12-13 minutes in the recovery chamber.

#### Suggestion

Start the timer after transferring the first slice and transfer them to the RT R-aCSF solution every 13 minutes to make the incubation time even. Too long incubation (>20 minutes) may make the neuronal membranes stiff and more difficult to harvest the somas.

27. After 12-13 minutes transfer the slices to the R-aCSF long-term holding chamber maintained at RT. Allow slices to recover for an additional 45-60 minutes in the holding chamber prior to initiating voltage imaging experiments.
28. **Incubation in JF dye:** Incubate slices at RT for 25-30 min in 50 nM Janelia-Fluor 585 dye dissolved in carbogenated R-aCSF. Transfer the JF-labeled slices back to the RT long-term holding chamber with R-aCSF or straight to the chamber of the electrophysiology setup.

#### Suggestion

29. In Our experience after 25-30 minutes the Halo-Tags are already saturated and longer incubation does not improve the signal-to-noise ratio.
30. It is possible to incubate all the slices together and transfer them back to the incubating R-aCSF until recording. The JF-dye binds to the Halo-Tag covalently, and there is no difference in intensity between slices imaged immediately after incubation or later in the day.

## All-optical voltage imaging

### Suggestion

First, place the slice with the injection site of the ChR2-expressing virus in the recording chamber and check the viral expression to validate the successful ChR2 expression, that is essential for the all-optical experimenting.

31. Place a slice with Voltron expression in the recording chamber and first validate the expression of ChR2 in the axons with low intensity 470-490 nm light and validate the Voltron expression with lowest possible intensity of 585 nm light.
32. Set the imaging FOV and set the focus wile using 585nm light with minimal/low intensity illumination to avoid bleaching of the JF-585.
33. In HCImageLive software, set the size of the FOV. We used 168 (y) x 480 (x) pixels. Width (x-axis) of FOV does not slow down frame acquisition.
34. While the LED light source is switched off, set both the intensity of the 473 nm blue light (for optogenetic stimulation) and the 585 nm yellow light (for voltage imaging) to the desired light power, in our case to ∼2.5 mW/mm2 and to ∼14 mW/mm2 respectively.
35. Make sure that exposure time is set to 1.65 ms, that will result in 600 Hz.
36. Check the “Frame bundle” box, that will concatenate multiple sweeps into one video file. Sweeps can be acquired as separate files, in this case the saving time may be longer then the intended intersweep-intervals. In line with this, concatenation will result in a longer saving time of even 2 minutes (depending on the frame rate and the length of the sweeps).
37. Set the trigger in the HCImageLive to “External Trigger” mode and start the experiment.
38. Start the protocol file In Axon 10.4. HCImageLive will only acquire the data when the camera receives the frame trigger coming from the Arduino with the speed of 600 Hz.
39. We acquired data with ∼10x magnification (20x objective with 0.5x demagnifier)
40. Run the On-site VoltView on the saved data (see VoltView_README.pdf at https://zenodo.org/record/8030176) and explore the postsynaptic responses in the VoltView ROI explorer. Here one can choose neurons based on user defined requirements for Voltage-Seq soma harvesting.

### Alternative experimental path

With the built-in on-site comparison of multiple videos taken from the same FOV (e.g., control and pharmacology), it is also possible to select neurons for harvesting based on the changes observed after a pharmacological treatment. Alternatively, data can be accumulated without on-site analysis and analyzed offline.

### Suggestion for 3D mapping

The cameras suitable for all-optical imaging often have nonstandard chip sizes, pixel size calibration is necessary and helpful for later mapping the recorded neurons.

## Voltage-seq soma harvesting

The harvesting protocol was optimized for Voltage-Seq, for scRNA-seq we used Smart-Seq2^10^.

### Suggestion

During the harvesting process RNAse free working environment is not necessary.

41. Fill the harvesting electrode with the smallest amount of harvesting solution possible.

### Suggestion

2-3 MΩ tapered pipettes are suitable to aspire the neuronal somas effectively. Pipette pullers used to create pipettes for whole-cell patch clamp experiments with 7-8 MΩ resistance. It can be a challenge to pull consistent 2-3 MΩ pipettes. Pull the harvesting pipettes the day before the experiment. Verify the resistance of the pipettes with a microforge (Narishige MF-900) or test every 4-5th pipette in the setup with the harvesting solution and with positive pressure.

### Suggestion

Filling the harvesting pipette can be standardized with a filler tip and pipette. After loading 1-2 ul harvesting solution, let the capillary effect of the tapered electrode fill in the thinnest part. With the help of the loading pipette tip take back the excess, what is left in the tapered part is enough to harvest.

42. Apply positive pressure to have a mild flow of harvesting solution. Approach the suggested soma with the harvesting pipette.
43. Aspire the entire soma of the selected neuron within a few seconds by applying mild negative pressure (−50 mPa) measured with a manometer.

### CRITICAL STEP

If the harvesting attempt takes minutes, it could mean the harvesting pipette is clogged and will not aspirate or eject the sample properly, resulting in low quality sample.

### Suggestion

Switching from DIC to fluorescent optics to visualize the Voltron-expressing neurons during and after aspiration could confirm the successful harvesting process.

44. Pull out the harvesting pipette of the recording chamber slowly.
45. With the micromanipulator, navigate over and inside a 0.2 ml tube under visual guidance of a lower magnification lens to validate sample ejection(preferably 4x air objective). Only dip the tip of the harvesting pipette into the LB, to keep the sample cleaner.
46. Apply positive pressure eject the harvested neuron. Through the 4X air objective we could observe the line of small air bubbles coming out of the harvesting pipette tip into the LB as a confirmation of the completed ejection. The resultant sample (∼4,5 μl) was spun down (15-20 s), placed to dry ice and stored at -80 °C, later subjected to in-tube reverse transcription.

### Suggestion

The LB contains SEQURNA thermostable RNase inhibitor, which remains effective at room temperature, so this step is not time sensitive.

47. **Cell lysis:** Thaw Voltage-seq samples (PCR tubes containing lysis buffer and collected neuron) on ice.
48. Incubate the samples at 72 °C for 3 minutes to perform cell lysis and oligo-dT hybridization to the mRNA. Immediately put the tubes back on ice.
49. **Reverse transcription:** Prepare the RT mix for all reactions plus several additional reactions by combining and mixing the reagents listed in the table below. (Ideally one will want to prepare the RT Mix while the samples are being lysed).

| Component | Final concentration | Volume (μl)<br>for 1 sample | Volume for 96<br>samples (x110) |
| --- | --- | --- | --- |
| SuperScript II reverse transcriptase (200 U μl<sup>-1</sup>) | 100 U | 0.50 | 55 |
| Superscript II first-strand buffer (5×) | 1× | 2.00 | 220 |
| DTT (100 mM) | 5 mM | 0.50 | 55 |
| Betaine (5 M) | 1 M | 2.00 | 220 |
| MgCl<sub>2</sub> (1 M) | 1 mM | 0.10 | 11 |
| Smart-seq2 TSO (100 μM) | 1 μM | 0.10 | 11 |
| Nuclease-free water | – | 0.30 | 33 |
| Total volume | – | 5.50 | 660 |

50. Add 5.5 μl of the RT mix to the samples from Step 32 to obtain a final reaction volume of 10 μl.
51. Spin down the samples (700*g* for 10 s at room temperature) to collect the liquid at the bottom of the tubes, and incubate the reaction in a thermal cycler with a heated lid, as described below:

| Cycle | Temperature (°C) | Time |
| --- | --- | --- |
| Step 1 | 42°C | 90 min |
| Step 2<br>10 cycles | 50°C | 2 min |
|  | 42°C | 2 min |
| Step 3 | 70°C | 15 min |
| Step 4 | 4°C | Hold |

52. **PCR preamplification:** Prepare the PCR mix for all reactions plus several additional reaction by combining and mixing the reagents from the table below. (Ideally you will want to prepare the Amplification Mix while your samples are near the end of reverse transcription).

| Component | Final concentration | Volume (µl) for 1 sample | Volume (µl) for 96 samples (x110) |
| --- | --- | --- | --- |
| KAPA HiFi HotStart ReadyMix (2×) | 1× | 12.5 | 1375 |
| ISPCR primers (10 µM) | 0.1 µM | 0.2 | 22 |
| Nuclease-free water | – | 2.3 | 253 |
| Total volume | – | 15 | 1650 |

53. Add 15 μl of PCR mix to each tube, which contains the first-strand reaction. Spin tubes down (700*g* for 10 s at room temperature) to collect the liquid at the bottom of the tubes.
54. Perform the PCR in a thermal cycler by using the following program:

| Cycle | Temperature (°C), Time | Process |
| --- | --- | --- |
| Step 1 | 98 °C, 3 min | Denature |
| Step 2<br>22 cycles | 98 °C, 20 s | Denature |
|  | 67 °C, 15 s | Anneal |
|  | 72 °C, 6 min | Extend |
| Step 3 | 72 °C, 5 min | Final Extension |
| Step 4 | 4 °C | Hold |

### Pause point

PCR product can be stored at −20 or −80 °C for 6 months or longer.

#### PCR purification

55. Prior to purification, allow 22% PEG Clean-up Beads (prepared as cited in dx.doi.org/10.17504/protocols.io.p9kdr4w) to equilibrate to room temperature for 15 minutes and mix thoroughly with vortexing to resuspend beads. These beads are equivalent to AMPure XP beads (Beckman Coulter).
56. Add 20 μl of 22% PEG Beads (0.8:1 beads:cDNA ratio) to each sample of pre-amplified cDNA and mix thoroughly to combine. Let incubate 8 minutes to bind DNA to beads.
57. Place on a compatible magnetic stand for 5 minutes or until the solution runs clear to collect the beads.
58. Keeping the samples on the magnet, remove the supernatant without disturbing the pellet.
59. Wash the beads while the samples are on the magnet with 200 μl of 80% (vol/vol) ethanol solution. Incubate the samples for 30 s and then remove the ethanol.
60. Repeat the ethanol wash once more.
61. Remove any residual ethanol and allow beads to dry completely, maximum 5 minutes or until a small crack appears on the surface of the beads.
62. Remove the sample from the magnet and add 17.5 μl of EB solution. Pipet up and down to thoroughly resuspend the beads.
63. Incubate the samples off the magnet for 2 minutes.
64. Place the samples on the magnetic for 2 minutes until the solution appears clear and beads have accumulated in a corner of the well.
65. Set the volume of the pipette to 15 μl, collect 15 μl of the supernatant and transfer it to a fresh PCR tube, 8-well PCR strip or 96-well plate.

#### Quality check of the cDNA library of Voltage-Seq samples

66. Check the yield and size distribution on an Agilent high-sensitivity DNA chip. A good library should be nearly free of short (<500 bp) fragments and should show a peak at ∼2 kb, corresponding to the median length of mRNAs in mouse and human cells^11^

#### cDNA quantification and normalization

67. We follow the QuantiFluor® dsDNA kit to measure concentration of samples.
68. Prepare standards as stated in the kit.
69. Make a working solution consisting of 1:400 dilution of QuantiFluor® dsDNA Dye in 1X TE buffer.
70. Add 49 μl per well to the 384 flat-bottom well well plates and 99 μL per well to the 96 well plates of the ready Quantiflour mix.
71. Add 1 μl of cDNA to each well (make sure you have enough extra wells to add 1 μl of the standards in duplicate). Incubate assay for 5 minutes at room temperature, shaking at 300 rpm.
72. Use a plate reader, to measure fluorescence (504nM Excitation/ 531nM Emission)
73. Calculate cDNA concentration and calculate water needed to dilute cDNA to between 0.2 ng to 1 ng/μl (ideally same concentration for every sample, if possible.
74. Prepare a new normalization plate by adding the calculated water volumes to each well.
75. Add 1-5 μl of preamplified cDNA to each well (depending on calculation).

#### Tagmentation reaction

Tagmentation is carried out by using an in-house Tn5, prepared according to Smart-Seq2^12^. (https://genome.cshlp.org/content/24/12/2033).

76. Set up the Tagmentation Mix on ice and pipette up and down carefully to mix:

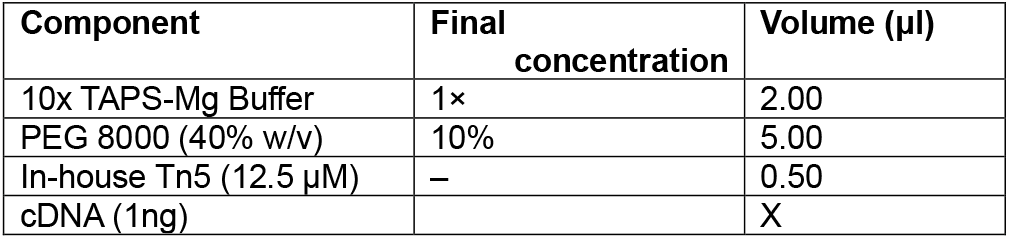
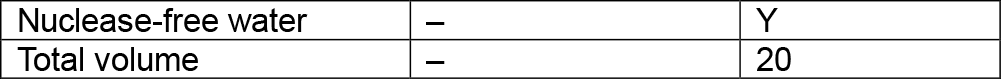

77. Perform the tagmentation reaction in a thermal cycler by using the following program:

| Cycle | Temperature (°C) | Time |
| --- | --- | --- |
| Step 1 | 55 | 5 min |
| Step2 | 4 | Hold |

#### Stripping Tn5 transposase off the tagmented DNA

78. To strip off the Tn5 bound to DNA, add 3.5 μl of 0.2% SDS to each sample well. Seal plate or close lid of tube and briefly vortex and spin down.
79. Incubate the mixture for 5 minutes at room temperature. DNA is now ready for the final enrichment PCR (indexing).

## Indexing

80. Add indexes to the tagmented DNA. Be careful not to cross contaminate indexes. If using index plates, be sure to spin down plates and gently pull back foil, replacing it with a *new* foil lid when done.

| Component |  | Volume (µl) |
| --- | --- | --- |
| Custom in-house library indexes |  | 1 |

81. Prepare the Library Amplification Mix on ice as follows. Final volume for the library amplification step should be **50 μl**, so water should be adjusted accordingly if you deviate from the SDS and primer volumes.

| Component | Volume (µl) | Final concentration |
| --- | --- | --- |
| Tagmented DNA + index | 24.5 | – |
| 5x HiFi PCR buffer | 10 | 1x |
| 10mM dNTP | 1.5 | 0.3 mM each |
| KAPA HiFi polymerase (1U/µl) | 1 | 1U |
| Molecular grade water | 13 | – |
| Total volume | 50 | – |

82. Next, perform the PCR by using the following program:

| Cycle | Temperature (°C), Time | Time |
| --- | --- | --- |
| Step 1 | 72 °C, 3 min | Pre-step |
| Step 2 | 95 °C, 30 s | Denature |
| Step 3<br>10 cycles | 95 °C, 10 s<br>55 °C, 30 s<br>72 °C, 30 s | Denature<br>Anneal<br>Extend |
| 4 | C, 5 min | Extension |
| 5 |  |  |

### Final library purification

83. Pool individual libraries into a 1.5 mL microcentrifuge tube. Take at least 5 μl from each sample.
84. Add 22% PEG Beads (0.8:1 beads:cDNA ratio) to the final library pool.
85. Place on a compatible magnetic stand for 5 minutes or until the solution runs clear to collect the beads.
86. Keeping the samples on the magnet, remove the supernatant without disturbing the pellet.
87. Wash the beads while the samples are on the magnet with 1 ml of 80% (vol/vol) ethanol solution. Incubate the samples for 30 seconds and then remove the ethanol.
88. Repeat the ethanol wash once more.
89. Remove any residual ethanol and allow beads to dry completely, maximum 5 minutes or until a small crack appears on the surface of the beads.
90. Remove the sample from the magnet and add 42.5 μl of EB solution. Pipet up and down to thoroughly resuspend the beads.
91. Incubate the samples off the magnet for 2 minutes.
92. Place the samples on the magnetic for 2 minutes until the solution appears clear and beads have accumulated in a corner of the well.
93. Set the volume of the pipette to 40 μl, collect 40 μl of the supernatant and transfer it to a 1.5 ml microcentrifuge tube.

## Quality and concentration check of the final library

94. Check the size distribution on an Agilent high-sensitivity DNA chip. Load the undiluted sample along with a 1:5 and 1:20 dilution of your library in case the sample is overloaded. As a rule of thumb, a broad peak with an average size of 250–1000 bp will be observed.
95. Use a Qubit fluorometer to quantify the library for the Illumina NextSeq library preparation kit (other sequencers and kits may be equally suitable).

## Library sequencing

96. Prepare the sequencing library using a NextSeq 500/550 library preparation kit – for the Voltage-seq samples we used a 150 cycle Nextseq 550 kit (paired end, 74bp reads). Aim to sequence to enough depth that you obtain at least 1 million reads per sample.
97. Run library on a NextSeq 550 next-generation sequencing instrument.

## Data processing

98. After the sequencing run, the output format (bcl) needs to be converted to fastq. This can be done with following command line for a 2x74 dual index. <u>Command:</u> bcl2fastq --use-bases-mask Y74N,I10,I10,Y47N --create-fastq-for-index-reads --no-lane-splitting -R /dir/sequencing_runs/project_run --output-dir /dir/projects/Voltage_seq/output

## Anticipated results

With successful viral expression of both the voltage sensor and the optical actuator at the targeted anatomical areas (Fig. 6a), the experiment has an average throughput of 1000-1500 all-optical voltage imaged postsynaptic neurons per animal. In a single FOV on average 30-80 postsynaptic neuronal connections are simultaneously probed (Fig. 6b). With the average signal-to-noise ratio, single APs, o-EPSPs, o-IPSPs, bursts and rebound activity can be captured and well reproduced across consecutive sweeps. All-optical voltage imaging with 6-7 sweeps in the same FOV can be repeated up to 6-7 times with comparable sensitivity (Fig. 6c). Above this number of repetitions, bleaching is going to increasingly influence the signal-to-noise ratio by first affecting only the subthreshold signals and above 10 repetitions affecting even the detection of APs.

**Fig 4.**
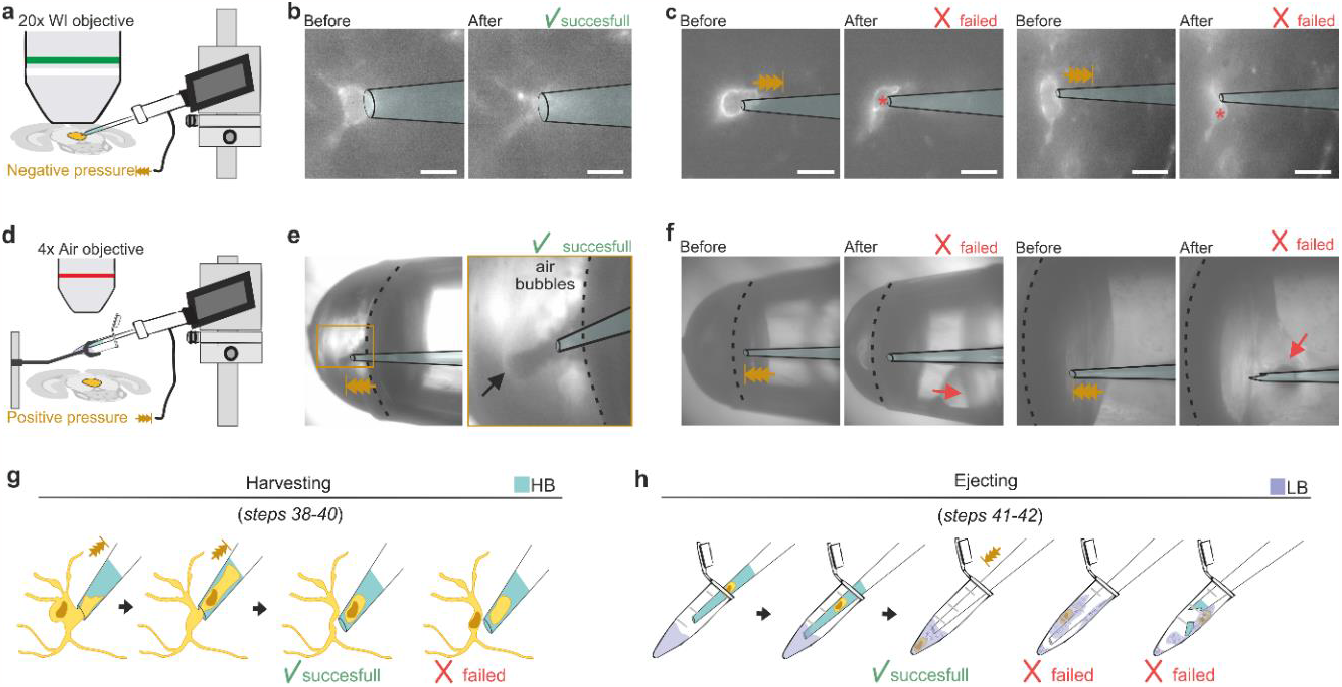
Voltage-Seq neuronal soma harvesting. **a**, Schematic of brain slice under 20x water immersion objective during harvesting with negative pressure. **b**, Successful harvesting of the entire soma. **c**, Two cases of failed soma harvesting due to high resistance harvesting capillary (>3,5 MOhm) where the nucleus (*) could not enter the small capillary tip. **d**, Schematic of harvesting PCR tube under 4x air objective with the harvesting stand positioned above brain slice. The harvesting stand holds the PCR tube with lysis buffer (LB) under the objective in fixed position. Harvesting capillary is navigated into the PCR tube with the micromanipulator and positive pressure is applied. **e**, Successful ejection of the harvested neuron with visible small bubbles in the LB indicating that the capillary is emptied. **f**, Failed ejection of the harvested somas, that typically result in low or no yield of cDNA. Ejection speed was excessively high, caused foaming up of LB that adhered to the wall of the PCR tube (left). Capillary broke against the wall of the PCR tube (right). **g**, Illustration outlining the neuronal harvesting procedure. Harvesting capillary contains the harvesting buffer (HB, blue) Successful harvesting includes the nucleus. Failed harvesting lacks the nucleus. **h**, Schematic illustration of the ejection of the harvested neuron to the lysis LB (purple). Successful ejection showing the formation of bubbles, and two cases of failed ejection due to high speed of ejection, and broken capillary.

**Fig 5.**
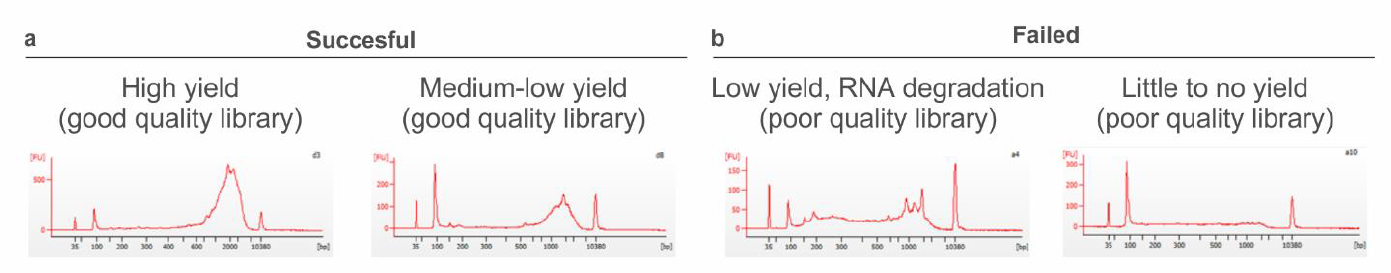
BioAnalyzer cDNA quality check. **a**, Successful Voltage-Seq cDNA libraries with high yield and medium-low yield. Below this quality the samples are considered to be discarded **b**, Failed cDNA libraries either because of RNA degradation or because the sample was almost empty. The latter occurs when the nucleus is not harvested or upon ejection into the harvesting PCR tube the sample foams up and that can lead to the RNA adhering to the wall of the plastic PCR tube.

**Fig 6.**
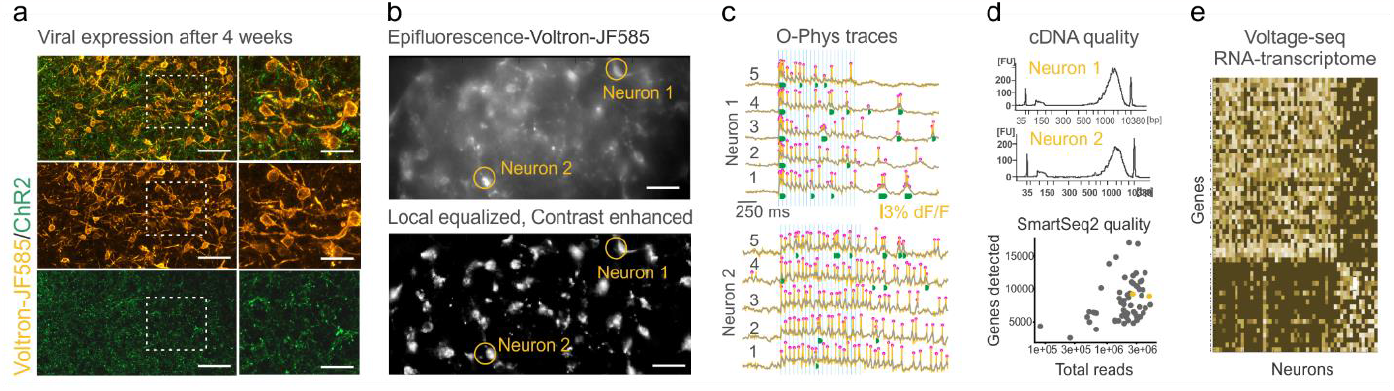
Anticipated results. **a**, Successful viral expression of Voltron-ST and ChR2 **b**, Example FOV in 1P epifluorescent optics with Voltron-JF-585 (top) and the preprocessed image after local equalization and contrast enhancement for soma segmentation. **c**, Example o-phys traces from the 2 neurons indicated in (b) with APs, subthreshold o-EPSPs, bursting activity and persistent AP firing well reproduced across consecutive sweeps. **d**, Example good-quality cDNA libraries from the same 2 neuron types after somatic harvesting and reverse transcription (top) and descriptive metrics of Smart-seq2 results. **e**, Differential expression analysis of excitatory vs. inhibitory neurons from our Voltage-Seq RNA transcriptome dataset generated in the original paper.

Using HCImageLive software generates a .cxd file, compatible with analysis with VoltView. Upon successful execution, VoltView generates a .mat file carrying the same name as the original video.

This .mat file contains the PARAMS file, described in greater detail in Table 2 and Table 3. The VoltView ROI Explorer runs in a 1-1.5 minutes and provides the PRTs of all-optical voltage-imaged neurons in the FOV.

By using Voltage-Seq, it is possible to collect 20-25 neurons a day. The quality check of the samples harvested without preceding whole-cell recording showed high or medium cDNA yield (Fig. 6d), indicating that the voltage sensor expression did not influence the overall quality of the neuronal mRNA. The descriptive metrics of the number of total reads and the number of detected genes per neuron after scRNA-seq are in line with the available open source scRNA-seq data (Fig. 6d). Analysis of the gene expression will reveal characteristics of the investigated neurons that allow to test for example peptide neuromodulation, as we demonstrated in the original paper (Fig. 6e).

